# TRIBE editing reveals specific mRNA targets of eIF4E-BP in *Drosophila* and in mammals

**DOI:** 10.1101/2020.02.24.962852

**Authors:** Hua Jin, Daxiang Na, Reazur Rahman, Weijin Xu, Allegra Fieldsend, Wei Song, Shaobo Liu, Chong Li, Michael Rosbash

## Abstract

4E-BP (eIF4E-BP) represses translation initiation by binding to the 5’cap-binding protein eIF4E and inhibiting its activity. Although 4E-BP has been shown to be important in growth control, stress response, cancer, neuronal activity and mammalian circadian rhythms, it is not understood how it preferentially represses a subset of mRNAs. We successfully used hyperTRIBE (Targets of RNA-binding proteins identified by editing) to identify in vivo 4E-BP mRNA targets in both *Drosophila* and mammals under conditions known to activate 4E-BP. The protein associates with specific mRNAs, and ribosome profiling data show that mTOR inhibition changes the translational efficiency of 4E-BP TRIBE targets compared to non-targets. In both systems, these targets have specific motifs and are enriched in translation-related pathways, which correlate well with the known activity of 4E-BP and suggest that it modulates the binding specificity of eIF4E and contributes to mTOR translational specificity.

## Introduction

InR-mTOR signaling is well conserved in all eukaryotes and integrates different environmental signals to govern organismal growth. Under favorable circumstances such as a nutrient-rich conditions, InR-mTOR signaling accelerates growth by increasing processes involved in anabolic metabolism and promoting growth in cell size and number (Saxton and Sabatini 2017). The same processes are inhibited in nutrient-restricted or stress conditions. A major process regulated by InR-mTOR signaling is translation. This is so that this high energy-consuming process can be increased under pro-growth conditions or decreased under low-growth or no-growth conditions, e.g., under stress conditions.

4E-BP (eIF4E-binding protein) is a crucial regulator of this translational control pathway. There are three homologs (4E-BP1, 4E-BP2 and 4E-BP3) in mammals but only one in *Drosophila melanogaster* (Thor/d4E-BP), the product of the *Thor* gene. d4E-BP is crucial for lifespan extension under oxidative stress and nutrient-restricted conditions (Zid et al., 2009), consistent with a prominent role in restricting growth. In fact, the impact of 4E-BP extends far beyond cellular stress management, e.g., it impacts cancer, proper neuronal activity and mammalian circadian entrainment (Teleman et al. 2005; Tettweiler et al. 2005; Mamane et al. 2006; Zid et al. 2009; Cao et al. 2013).

4E-BP inhibits translation initiation by blocking the formation of the translation initiation complex eIF4F (eukaryotic initiation factor 4F) (Ma and Blenis 2009; Kong and Lasko 2012; Saxton and Sabatini 2017). eIF4F consists of the 5’cap-binding protein eIF4E, the RNA helicase eIF4A and the scaffold protein eIF4G. Under favorable growth conditions, the mTOR signaling pathway phosphorylates 4E-BP, which prevents it from binding to eIF4E. Under less favorable or stress conditions, inactivation of mTOR signaling results in dephosphorylation and 4E-BP activation. Because eIF4G and hypophosphorylated 4E-BP share the same binding site on eIF4E, active 4E-BP blocks the eIF4G binding site on eIF4E and thereby inhibits eIF4F formation and translation.

To identify mRNAs regulated by 4E-BP, others have examined global mRNA translation using ribosome profiling under mTOR inhibition conditions (Hsieh et al. 2012; Thoreen et al. 2012). The results showed that active 4E-BP preferentially represses translation of a subset of mRNAs. Although inhibition of eIF4F formation can explain how the general rate of cap-dependent translation is decreased, it does not explain how the translation of specific mRNAs is selectively inhibited, i.e., whether this is a direct or indirect effect of active 4E-BP.

In vitro assays have suggested that 4E-BP may indirectly associate with mRNA through the cap binding protein eIF4E. There are several pieces of evidence supporting this hypothesis, including that 4E-BP can be co-purified with eIF4E from cell extracts with a mRNA-cap analog pulldown assay and that 4E-BP enhances the in vitro interaction of eIF4E with a capped short RNA oligonucleotide (Pause et al. 1994; Ptushkina et al. 1999). However, these interactions do not address the specificity issue, and it is uncertain whether they are physiologically relevant. This is because there are few tools to detect the association of proteins like 4E-BP with specific mRNA within cells.

To address this deficiency, i.e. to identify the in vivo target transcripts of RNA-binding proteins (RBPs) in *Drosophila*, we recently developed TRIBE, Targets of RNA-binding proteins identified by editing (McMahon et al. 2016). This technique characterizes fusion proteins between the catalytic domain of the *Drosophila* RNA editing enzyme ADAR (ADARcd) and RBPs of interest. A RBP-ADARcd fusion protein can bind to target transcripts of the RBP and edit adenosine nucleotides to inosines near the fusion protein binding site; these are read as guanosines by high-throughput sequencing and identified as A-to-G editing sites by computational analysis. Transcripts hosting these A-to-G editing sites are considered binding targets of the RBP. The upgraded version of TRIBE, hyperTRIBE, contains the one amino acid substitution E488Q in the ADARcd region (hyper-ADARcd), which improves target identification efficiency by reducing bias originating from the intrinsic editing preference of the ADARcd (Kuttan and Bass 2012; Rahman et al. 2018; Xu et al. 2018).

Here we applied hyperTRIBE to both flies and mammals and characterized 4E-BP direct target transcripts within cells. These transcripts were enriched for mRNAs encoding translation components, including the translation initiation factor eIF3. Translation factors fit with previous results describing 4E-BP-regulated transcripts, suggesting that much of this regulation is due to a direct interaction of 4E-BP-containing protein complexes with target transcripts.

## Results

mTOR-dependent translational repression affects specific genes, suggesting that 4E-BP may associate with a subset of mRNAs and/or that some mRNAs selectively escape from 4E-BP-mediated repression (Marr et al. 2007; Thoreen et al. 2012). The TRIBE method was developed in the *Drosophila* system and is ideal to address the possibility that d4E-BP associates with specific mRNAs. To perform Thor-TRIBE experiments in cultured *Drosophila* S2 cells, we constructed a plasmid in which the hyperactive *Drosophila* ADARcd (hyper-dADARcd) was fused to *Drosophila* 4E-BP (d4E-BP) (Suppl. Fig. 1A). If d4E-BP associates with specific eIF4E-5’capped mRNAs in cells, the hyper-dADARcd should deaminate nearby adenosines, leaving A-to-I editing marks on the associated mRNAs. A plasmid that only expresses the hyper-dADARcd was used as a negative control (Xu et al. 2018). We used western blotting to confirm their similar expression (Suppl. Fig. 1B) and also that Thor-hyper-dADARcd could be efficiently dephosphorylated after mTOR inhibition (Suppl. Fig. 1C).

To address whether TRIBE can identify Thor-associated mRNA targets, mRNA libraries were generated from positively-transfected cells and sequenced with the Illumina Nextseq 500 System. To generate lists of editing sites and target genes, the sequencing data were then analyzed with our published TRIBE computational analysis pipeline (Rahman et al., 2018). In wild type S2 cells or S2 cells expressing hyper-dADARcd alone, very few editing events were detected. However, thousands of editing sites were detected after induction of Thor-TRIBE expression (Fig. 1A, Thor-Hyper). Since mTOR inhibition dephosphorylates 4E-BP and enhances its activity and its interaction with eIF4E (Thoreen et al. 2012), we reasoned that mTOR inhibition should increase the number of target mRNAs identified by Thor-TRIBE. Indeed, more editing sites and target genes were reproducibly detected after inhibiting mTOR activity with serum depletion and rapamycin treatment. We also carried out Thor TRIBE experiments after rapamycin or torin treatment, which also inhibit mTOR activity. The data taken together indicate that Thor-TRIBE reflects d4E-BP activity and that the fusion protein associates with a subset of specific mRNAs in cells (Fig. 1A).

**Fig. 1.**
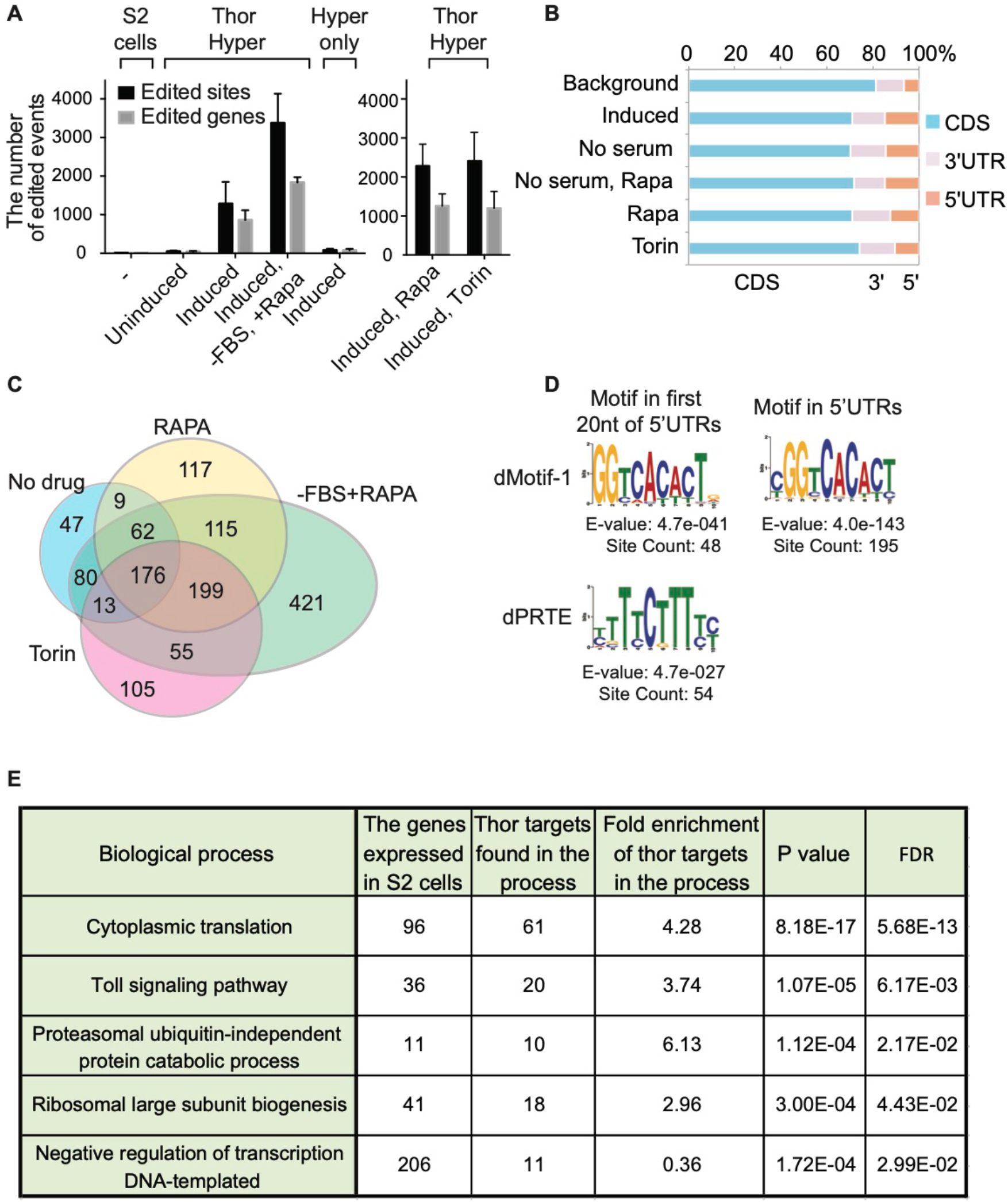
Thor hyperTRIBE identifies specific RNAs as 4E-BP targets in *Drosophila* S2 cells. S2 cells were transfected with a plasmid that contains Thor-hyperADARcd (Thor Hyper) or hyperADARcd alone (Hyper only) under copper inducible metallothionein (MT) promoter. TRIBE construct expression was uninduced or induced by 0.5 mM CuSO_4_ for 16 hours. For mTOR inhibition, cells were incubated with 50 nM of Rapamycin or Torin for 5 hours before harvesting. **(A)** Thor hyperTRIBE but not Hyper only edits transcripts after copper induction. The editing sites (black bars) and editing genes (gray bars) in Thor hyperTRIBE significantly increased in Rapamycin or Torin treated cells and more than doubled in Rapamycin treated plus serum deprived cells. The number of editing events in cells expressing hyperADARcd alone is comparable to that of control S2 cells. (N=2, +SEM) **(B)** Editing sites identified by Thor hyperTRIBE are enriched in 5’UTR of mRNAs. Background distribution was calculated by read distribution tool in RSeQC package, ~80% of the sequencing coverage was in coding sequencing (CDS, blue), ~15% in 3’UTR (green) and ~5% in 5’UTR (purple). Editing site distribution was calculated by Bedtools intersect with dm3 Ref-seq annotation. **(C)** Venn diagram of Thor hyperTRIBE target RNAs reproducibly identified under different conditions shows that the targets are consistent, although significantly more were identified with serum deprivation and Rapamycin treatment combined. The transcripts edited in S2 or Hyper only control were removed from target list. 176 targets genes were identified in all conditions. **(D)** The consensus motifs from the 5’UTRs of 968 Thor hyperTRIBE targets (listed in Suppl. Table 1) which were reproducibly detected in Rapamycin or Torin treatment condition are shown. These motifs were not detected from randomly-selected same number of non-target genes. The motifs from first 20 nt of 5’UTRs in Thor hyperTRIBE targets are shown on the left and the motif from entire 5’UTRs of targets is shown on the right. The GGTCACACT motif is identified in both cases with 195 counts (~20=) in entire 5’UTRs of the targets. The pyrimidine rich motif (dPRTE) resembles the PRTE motif previously implicated with mTOR-regulated RNAs (Hsieh et al. 2012). **(E)** Table of enriched GO term biological processes in 968 Thor hyperTRIBE targets reproducibly detected in Rapamycin or Torin treatment condition are shown. Genes involved in translation process and Toll signaling pathway are enriched in the target list. Fold enrichment > 1 means that targets are enriched in the process and fold enrichment < 1 means that targets are depleted in the process.

We next performed metagene analysis to examine the distribution of editing sites. The distribution of sequencing reads from all exonic regions was used as background. The editing sites of Thor-TRIBE were relatively enriched in 5’UTR regions compared to this background, consistent with a preferred association of d4E-BP with the eIF4E-5’cap complex (Fig. 1B). The target transcripts identified under these different conditions overlapped well, although many more were detected after mTOR inhibition (Fig. 1C). We carried out de novo motif searches from the 5’UTRs or 3’UTRs of the 968 Thor targets (Suppl. table 1), which were reproducibly detected in Thor-hyperTRIBE after Rapamycin or Torin treatment. As a background control, we randomly selected 968 expressed genes, which were not identified as Thor targets.

The search for consensus sequences from the first 20nt of target 5’UTRs identified a GGTCACACTG motif (dMotif-1) and a Pyrimidine-Rich motif (dPRTE, Fig. 1D). A further motif search within the full 5’UTR of targets identified a similar CGGTCACACT motif with an even more significant E-value and site count than that from the first 20nt; no significant motifs were found within the 3’UTR.

The dPRTE resembles the 5’terminal oligopyrimidine tract (5’ TOP) and the Pyrimidine Rich Translational Element (PRTE) identified as translational control elements in mammals (Suppl. Fig. 1D)(Meyuhas 2000; Tang et al. 2001; Meyuhas and Dreazen 2009; Hsieh et al. 2012). These motifs are also enriched within the 5’UTRs of mTOR-responsive mammalian mRNAs (Hsieh et al. 2012; Thoreen et al. 2012), consistent with the results shown here. Both the 5’TOP and the PRTE are stretches of uninterrupted pyrimidine nucleotides, but the 5’TOP starts with a cytosine residing from position +1 of the 5’UTR, and the PRTE is defined as a pyrimidine-rich element within the 5’UTR including an invariant uridine at position 6 of the motif. The activity of mammalian LARP is also under mTOR regulation and is reported to bind to a similar 5’TOP or PRTE motif (see Discussion)(Hong et al. 2017; Lahr et al. 2017).

Furthermore, GO term analysis indicates that these Thor targets are enriched in protein synthesis pathways, consistent with notion that translation is a key process regulated directly by Thor (Fig. 1E). Toll signaling and ubiquitin-independent proteasomal proteins are also enriched, whereas negative regulators of transcription are depleted in the Thor targets. As the mTOR pathway has been shown to be an important mediator of immune responses (Powell et al. 2012; Weichhart et al. 2015), these results implicate Thor in regulating the innate immune response via Toll signaling.

To verify that d4E-BP inhibits the translation of these mRNAs, protein synthesis was assayed in a number of different ways. We first used metabolic labeling by SUnSET to assay the effect of mTOR inhibition on general translation. This method utilizes puromycin incorporation into newly synthesized peptides to monitor the rate of general protein synthesis (Schmidt et al. 2009). Puromycin-attached peptides are detected by western blotting using an anti-puromycin antibody. Cells were pulsed with puromycin, and we compared general protein synthesis after treating cells with several mTOR inhibitors: Rapamycin, Torin, or Ink128. As expected, all three inhibitors reduced protein synthesis, with Rapamycin the most effective (Fig. 2A).

**Fig. 2.**
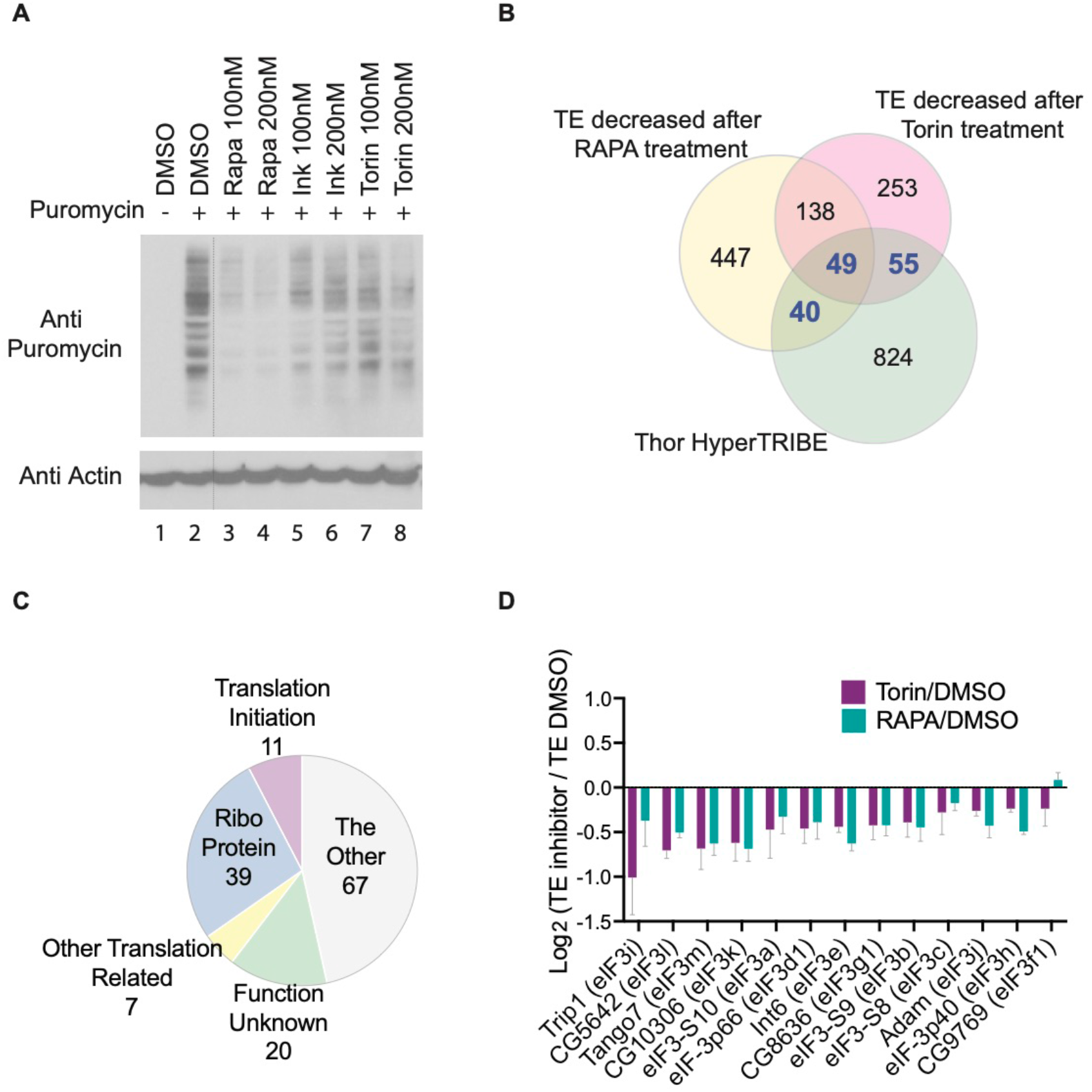
Ribosome profiling in S2 cells identifies mRNAs regulated by mTOR pathway. **(A)** Metabolic labeling by SUnSET shows global protein synthesis reduced after treating S2 cells with mTOR inhibitor, Rapamycin, Ink128 or Torin for 2 hours compared to vehicle DMSO. Puromycin was incorporated into newly synthesized peptides, being measured by Western blotting using antibody against puromycin. **(B)** Ribosome profiling identifies 144 Thor TRIBE targets (40+49+55, gene numbers are marked by blue color) translational efficiency (TE) of which decreased after treating cells with 100 nM of Rapamycin or Torin. Venn diagram overlap of genes between 968 Thor TRIBE targets, genes with decreased TE after adding Rapamycin, and genes with decreased TE after adding Torin. (n=3, TE fold change of mTOR inhibitor versus DMSO<0.8, p<0.1) **(C)** Functional classification of 144 Thor TRIBE targets which are from the overlapped portion in (B). The main functions of these genes include Ribosome components (Ribo Protein), translational initiation, and other translation-related roles. **(D)** The translation of most eIF3 mRNAs is repressed by mTOR inhibitor. The bar chart showed log_2_ TE-fold-change value of all expressed eIF3 transcripts. (Rapamycin/DMSO blue bar, Torin/DMSO purple bar, n=3, +SEM)

To examine translational activity at the whole transcriptome level, we carried out ribosome profiling with and without mTOR inhibition, i.e., 100nM of Rapamycin or Torin for 2 hours; the former should have a stronger effect and the latter a weaker effect on general translation. The strategy of ribosome profiling in mammals and yeast is to obtain ribosome-protected fragments (RPFs) by digesting cell lysates with RNase I (Ingolia et al. 2009; Ingolia et al. 2011). However, *Drosophila* ribosomes are reported to be too sensitive to RNase I, so micrococcal nuclease (MNase) is commonly used for ribosome profiling in this species. Although this makes P-site mapping difficult due to the strong 3′ A/T bias of MNase, it is still useful for measuring translation rates (Dunn et al. 2013).

Treating cell lysates with MNase confirmed that RPFs are enriched between 28nt and 34nt in size (Suppl. Fig. 2). We then purified these RPFs, made sequencing libraries using the adaptor ligation-based method and sequenced them. Consistent with the results of the metabolic labeling (Fig. 2A), more translationally-repressed genes were detected after treating with Rapamycin than that with Torin, i.e., 674 mRNAs decreased in translational efficiency (TE) after Rapamycin treatment and 495 mRNAs decreased in TE after Torin treatment (Fig. 2B and Suppl. table 2).

To orthogonally validate the Thor-TRIBE target data, we compared these genes with those that manifest decreased translational efficiencies in response to Rapamycin or Torin treatment. 144 of these transcripts were also identified by Thor-TRIBE, indicating that they are direct targets of d4E-BP (40 + 49 + 55; Fig. 2B). They encode ribosomal proteins, translational initiation factors, and other translation related proteins (Fig. 2C). Interestingly, these transcripts are highly enriched for members of the eukaryotic initiation factor 3 (eIF3) family. Of the 13 expressed members of this family, 10 fall in this overlapping category, namely eIF3b, eIF3d1, eIF3e, eIF3g1, eIF3h, eIF3i, eIF3j, eIF3k, eIF3l and eIF3m (Suppl. table 3). The TE of these 10 genes meaningfully decreased after mTOR inhibition (Fig. 2D).

We next asked whether the motifs enriched in Thor TRIBE targets play roles in translational repression. We compared the mean TE change after mTOR inhibitor treatment among different groups of transcripts expressed in S2 cells, including non-Thor TRIBE targets, all Thor TRIBE targets, Thor TRIBE targets with dMotif-1 or Thor TRIBE targets with dPRTE (Fig. 3A).

**Fig. 3.**
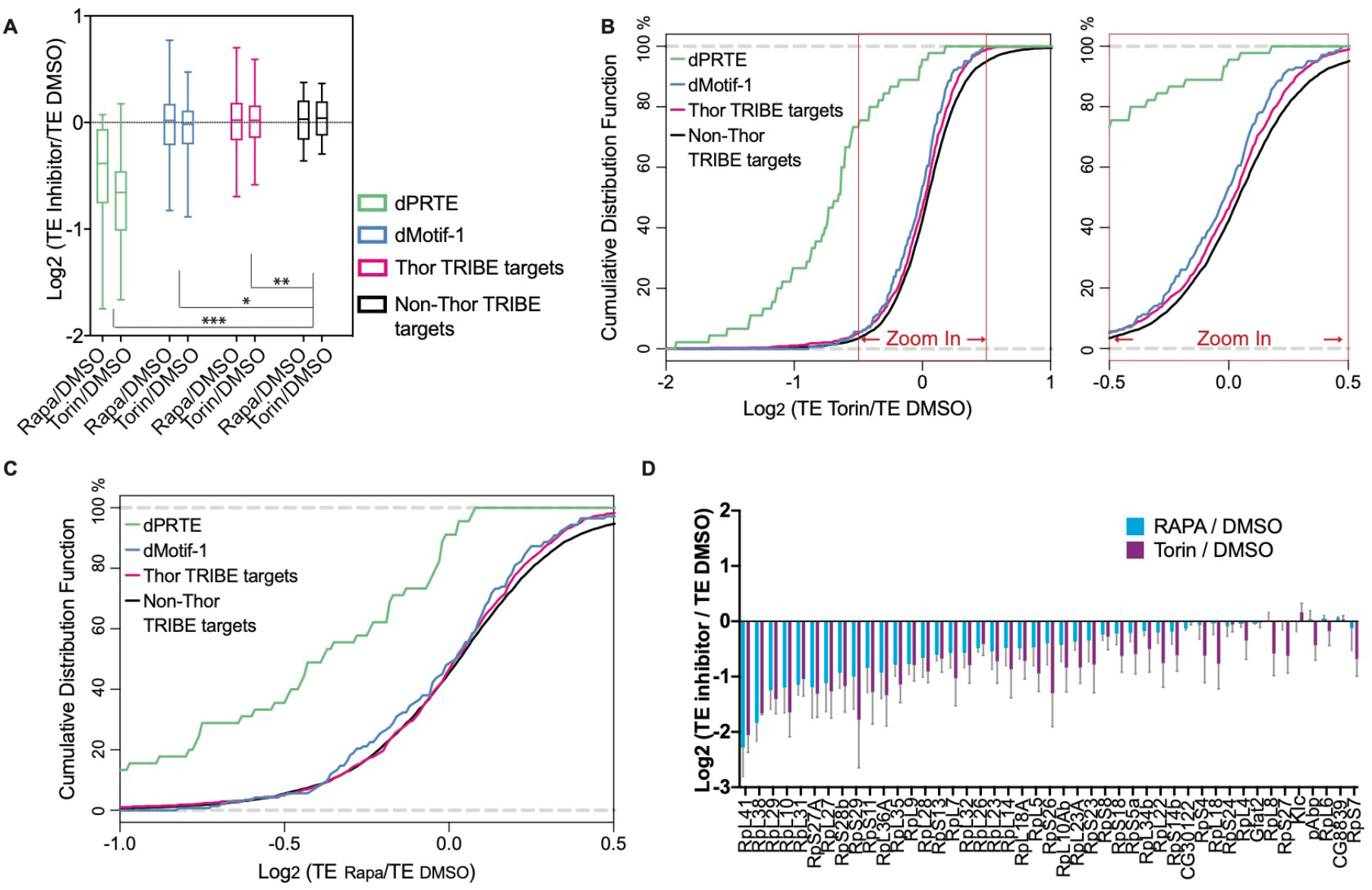
Ribosome profiling data show that the TE change of Thor TRIBE target after mTOR inhibition has a down-regulation compared with non-Thor TRIBE target. **(A)** The mean translational efficiency (TE) change of Thor TRIBE targets is down regulated comparing with non-targets when mTOR pathway is inhibited by mTOR inhibitor, Rapamycin or Torin. The degree of decrease varies with different groups of Thor TRIBE targets. Box plot of mean TE change (Log_2_ value) after a 2-h treatment with mTOR inhibitor versus vehicle DMSO from different groups of expressed transcripts, including all non-Thor TRIBE targets (black, n=5604), Thor TRIBE targets (purple, n=919), Thor TRIBE targets with dMotif-1 (blue, n=142) and with dPRTE (green, n=45). (Ordinary one-way ANOVA. * p<0.05, ** p<0.01, *** p<0.001) **(B-C)** Cumulative frequency distribution of log_2_ TE-fold-change values from non-Thor TRIBE targets (black, n=5604) and different groups of Thor TRIBE targets showed that Thor targets have significantly different TE change from non-targets. Torin vs. DMSO shown in (B); Rapamycin vs. DMSO shown in (C). **(D)** The TE of most Thor TRIBE targets with dPRTE decreased after treatment of mTOR inhibitor. The bar chart showed log_2_ TE-fold-change value of Thor TRIBE targets with dPRTE. (Rapamycin/DMSO blue bar, Torin/DMSO purple bar, n=3, +SEM)

Thor TRIBE targets containing the dPRTE/TOP motif showed dramatic translational repression in response to mTOR inhibition. This is consistent with previous reports indicating that that mTORC1 regulation of TOP mRNA translation is 4E-BP-dependent ((Hsieh et al. 2012; Thoreen et al. 2012; Nandagopal and Roux 2015)). Although less striking, the mean TE change of all Thor-TRIBE targets or of targets with dMotif-1 was also significantly different from that of non-targets. Cumulative frequency distributions of TE changes of targets with a dPRTE motif also had a striking distribution difference compared to non-targets, and all Thor TRIBE targets or targets with dMotif-1 showed a weaker but still significant difference from non-targets, especially when Torin was used as the mTOR inhibitor (Fig. 3B and Fig. 3C). The ribosome profiling data are notable because most dPRTE-containing Thor TRIBE targets had decreased TE after treatment with the mTOR inhibitors comparing to the DMSO control (Fig. 3D). Although the 5’TOP and the PRTE cis-regulatory motifs are well known in mammals and predicted to be translation elements in *Drosophila* (Suppl. Fig. 1D) (Meyuhas and Dreazen 2009), our results confirm that they play a conserved role in flies (Fig. 3D).

We then used CLIP to address RNA recognition and ask whether d4E-BP is in physical contact with RNA. The RBP and its UV-crosslinked RNAs in the cell lysate were partially digested with RNase A. After immunoprecipitating d4E-BP with a V5 antibody, crosslinked RNAs in the RBP were labeled with γ-[32P] ATP at their 5’ends and detected on denaturing polyacrylamide gels (Fig. 4A). A parallel control experiment was carried out with untransfected S2 cells and an antibody directed against the *Drosophila* HnRNP protein Hrp48. Although the Thor-TRIBE signal was not as strong as a typical RBP like Hrp48, radioactive signals were reproducibly detected by phosphoimager at 5-10kDa above the expected protein size, indicating that d4E-BP is in close proximity to RNA and may contribute to RNA binding activity like Hrp48 (Fig. 4B). Notably, transcript specificity of the CLIP tags was poor compared to the TRIBE data (data not shown), suggesting that Thor interacts broadly with mRNA and that TRIBE may be more successful at pointing to a biologically meaningful subset of transcripts (see Discussion).

**Fig. 4.**
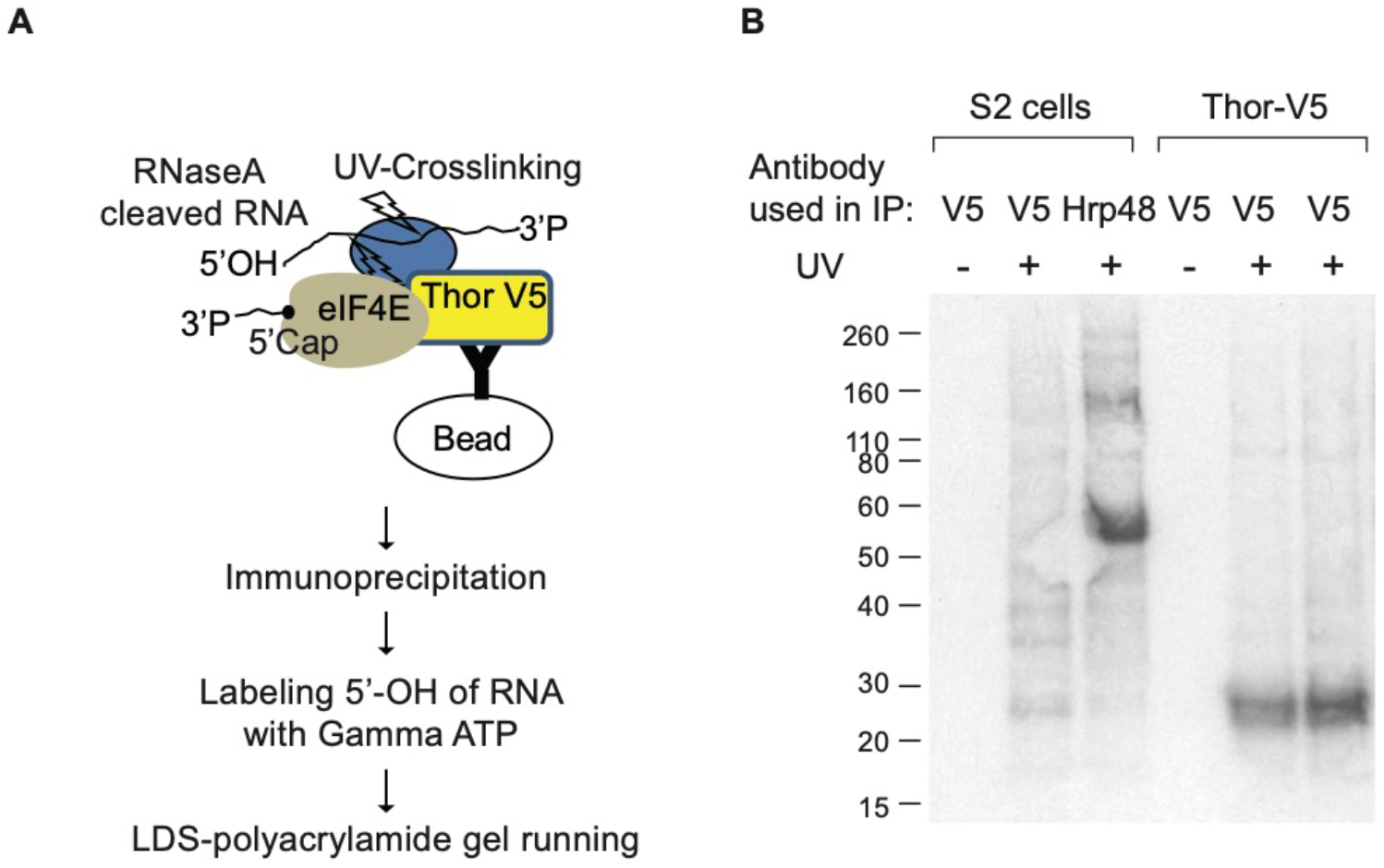
CLIP of Thor-V5 identifies a direct RNA-binding activity. **(A)** The schematic illustration of crosslinking-immunoprecipitation (CLIP). UV-crosslinked and lysed Thor-V5 cells were treated with RNase A, then immunoprecipitation was carried out from cell lysate using an antibody against V5 tag. The partially digested RNAs which had been crosslinked to RNA-binding protein (RBP) were labeled at their 5’end by γ-[32P] ATP. Radioisotope-labelled RNA-RBP could be separated and visualized in denaturing LDS polyacrylamide gel. **(B)** CLIP of Thor-V5 shows Thor has a RNA-binding activity. A well-known RBP, Hrp48 was used as a positive control. S2 cells or Thor-V5 cells were UV-crosslinked (UV+) or not (UV−). Immunoprecipitation was carried out using antibody against V5 tag or Hrp48.

Lastly, we examined whether 4E-BP also associates with target mRNAs in mammalian cells. As mentioned above, there are three 4E-BP proteins in mammals, namely, 4E-BP1, 4E-BP2 and 4E-BP3. They have somewhat different expression patterns in adult mouse tissues, and we chose 4E-BP1 (human or h4E-BP1) for hyperTRIBE experiments because it is the most broadly expressed (Tsukiyama-Kohara et al. 2001). To carry out h4E-BP1-hyperTRIBE in human cells, we first tried the hyperactive *Drosophila* ADARcd previously used for hyperTRIBE editing (Xu et al. 2018). However, few editing events were detected when *Drosophila* hyperADARcd was fused to h4E-BP1 and hyperTRIBE carried out in human prostate cancer cell PC3 cells (data not shown). We therefore substituted the hyper-dADARcd with the catalytic domain of human ADAR2 containing the E488Q point mutation at the corresponding position (hyper-hADAR2cd).

The results showed that h4E-BP1-hyperTRIBE but not the hyper-hADAR2cd alone successfully identified h4E-BP1 targets in PC3 cells. mTOR inhibition with Ink128/PP242 increased h4E-BP1-hyperTRIBE editing compared to the DMSO control (Fig. 5A), similar to the *Drosophila* results shown above (Fig. 1A). Also similar to those results, these human cell data indicate that 4E-BP inhibits translation by associating with its target mRNAs.

**Fig. 5.**
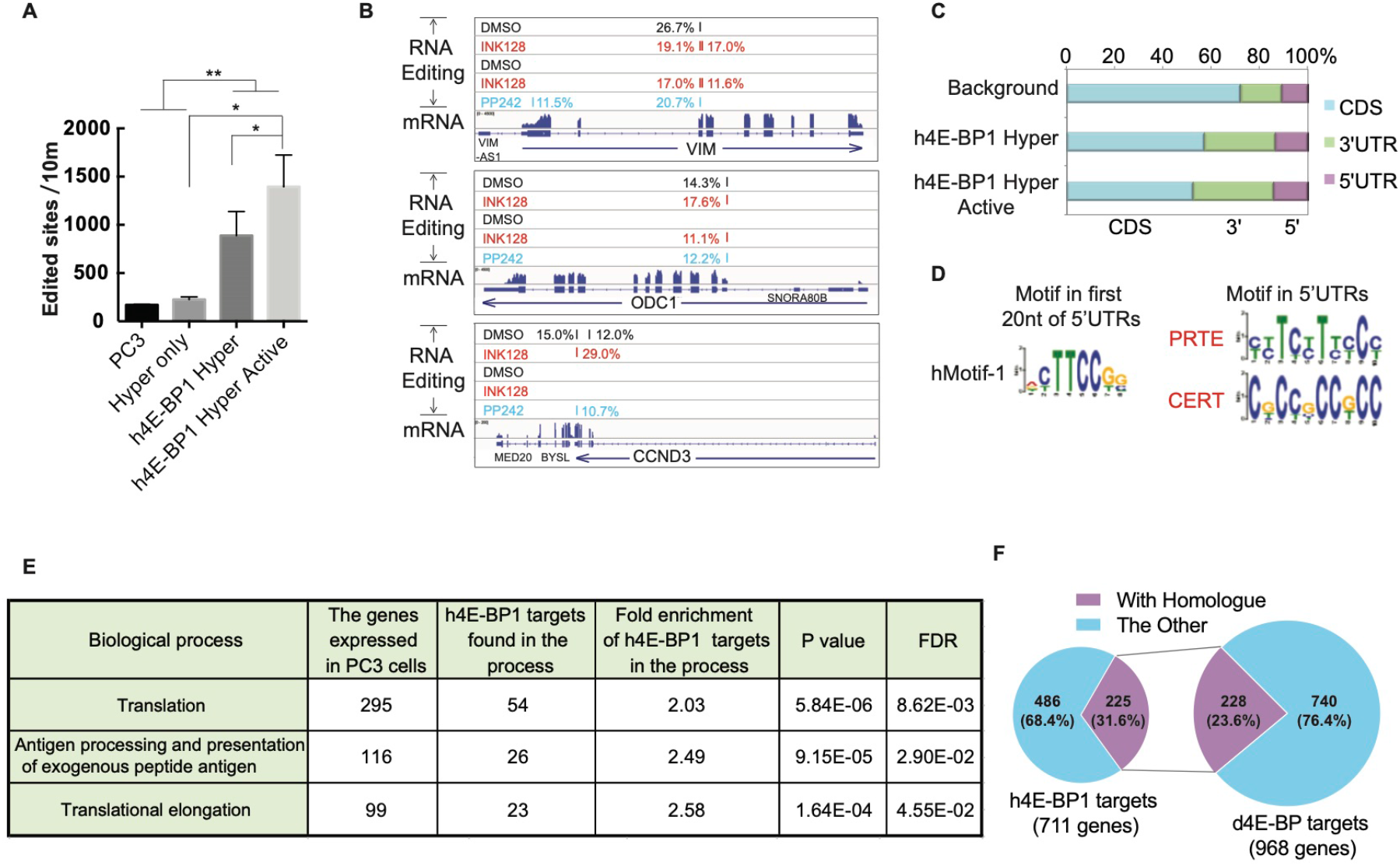
h4E-BP1 hyperTRIBE identifies h4E-BP1 targets in PC3 cells. PC3 cells were co-transfected a eGFP-expressing plasmid with a plasmid that contains h4E-BP1-hyperADARcd (h4E-BP1 hyper) or hyperADARcd alone (Hyper only) under CMV promoter for 16~40 hours. For mTOR inhibition, PC3 cells were incubated with 200 nM of INK128 or 2.5 μM of PP242 for 5 hours before harvesting. eGFP-positive cells were sorted by FACS and collected for mRNA-seq library preparation. **(A)** The number of edited sites is significantly higher in h4E-BP1 hyperTRIBE than that in wild type PC3 cells or in Hyper only. The number of edited sites was increased after INK128 or PP242 treatment (h4E-BP1 hyper Active). Expressing hyperADARcd alone (Hyper only) did not result in more editing sites than in control PC3 cells. (N=2~3, +SEM, * p<0.05 paired one-tail student TTEST, ** p<0.005 unpaired one-tail student TTEST) **(B)** IGV view of three examples of the editing sites and editing percentages of mTOR-sensitive genes VIM, ODC1 and CCND3 are shown. Background editing sites detected in control PC3 cells or Hyper only were removed. Samples were incubated with DMSO (black color), INK128 (red color), or PP242 (blue color). The alignment of mRNA-seq reads is also shown. **(C)** The editing sites of h4E-BP1 hyperTRIBE are enriched in 5’UTR and 3’UTR of mRNA. Editing site distribution and background coverage were calculated as described in Fig. 1B. **(D)** The consensus motifs from the 5’UTRs of 711 h4E-BP hyperTRIBE targets (listed in Suppl. Table 4) which were detected at least twice after mTOR inhibition. These motifs were not detected from randomly-selected same number of non-target genes. The motifs from first 20 nt of 5’UTRs in h4E-BP1 hyperTRIBE targets are shown on the left and the motifs from entire 5’UTRs of the targets are shown on the right. **(E)** Table of enriched GO term biological processes in 711 h4E-BP1 hyperTRIBE targets are shown. Genes involved in translation process and adaptive immune response are enriched in the target list. **(F)** Ortholog mapping between 711 h4E-BP1 hyperTRIBE targets and 968 Thor hyperTRIBE targets shows that 32% of h4E-BP1 targets are conserved in *Drosophila*. The purple fraction represents the shared 4E-BP targets that are orthologous in two species.

Transcripts of Vim (vimentin), ODC1 (ornithine decarboxylase) and CCND3 (cyclin D3) were previously identified as 4E-BP targets by polysome analysis or ribosome profiling using 4E-BP1/2 double knock-out (DKO) mouse embryonic fibroblasts (MEFs) (Dowling et al. 2010; Hsieh et al. 2012). These mRNAs were specifically edited by h4E-BP1-hyperTRIBE (Fig. 5B). Metagene analysis showed that these edited sites in h4E-BP1-hyperTRIBE were enriched in 5’UTR and 3’UTR sequences (Fig. 5C).

De-novo motif searches from the UTR regions of TRIBE targets identified the hMotif-1 in first 20nt of 5’UTRs. The PRTE as well as cytosine-enriched regulator of translation (CERT) were also identified within the entire 5’UTR region. The CERT motif was previously reported to be enriched in eIF4E-sensitive mRNAs, i.e., the translation of these transcripts was sensitive to eIF4E expression levels (Truitt et al. 2015).

These PC3 cell experiments identified 711 h4E-BP1 target genes (Suppl. table 4), and their functions were enriched in translation processes and the immune response like in fly cells (Fig. 5E). Comparing these 711 h4E-BP1 targets in human cells with the 968 d4E-BP targets in fly cells identified 180 sets of targeted homologs. As 225 human genes have 228 orthologues in flies, at least 32% of the 4E-BP human targets are conserved in flies (Fig. 5F and Suppl. table 5). These data point to conserved 4E-BP functions and mechanisms, and they also indicate that TRIBE works well in mammalian systems as well as in *Drosophila*.

## Discussion

TRIBE was first developed in *Drosophila* and worked well both in S2 tissue culture cells and in fly neurons (McMahon et al. 2016; Luo et al. 2018; Xu et al. 2018). To adapt this method to mammalian systems, we first tried the improved hyperTRIBE version with its hyperactive form of the *Drosophila* ADAR catalytic domain. For unknown reasons however, its editing efficiency was weak in PC3 cells (data not shown). So we turned to hyper-ADARcd from human ADAR2, which worked well in this human prostate cancer cell line.

The results indicate that 4E-BP associates in both systems with overlapping sets of target mRNAs, which are highly enriched in translation and immune response transcripts. TRIBE therefore captures the two major roles of mTOR, growth control and the immune response. Specific targets include translation factor mRNAs like those encoding ribosomal proteins (RPs); ribosomes are major growth effectors. In addition, 4E-BP represses the translation of most family members of the translation initiation factor eIF3, although to a lesser extent than RPs. The function of these targets is somewhat less straightforward to interpret, but one possibility is that these interactions also serve to inhibit growth: in this case reducing eIF3 protein levels would inhibit the translation of eIF3-dependent transcripts, which are less directly dependent on the canonical eIF4E-cap interaction (Lee et al. 2016).

To our knowledge, there are no previous data describing in vivo 4E-BP target transcripts. The association of 4E-BP with target RNAs increased after mTOR inhibition, conditions under which 4E-BP is reported to maintain a stronger interaction with eIF4E. It is interesting that Rapamycin has a stronger inhibitory effect on general ribosome profiling than Torin, whereas Torin has a stronger effect on the inhibition of mRNAs containing the specific motifs (Figs. 2 and 3). This suggests that the two inhibitors work somewhat differently, with Torin affecting more directly 4E-BP-4E-RNA complex formation.

Although previous results indicated that a subset of mRNAs is preferentially affected by mTOR and suggested that 4E-BP is the principle downstream translational repressor (Hsieh et al. 2012; Thoreen et al. 2012), it was unclear how mRNA specificity is determined. It had been proposed that expression of eIF4E is rate-limiting under some conditions such as tumorigenesis and that the sequestration of eIF4E by activated 4E-BP therefore reduces the level of the translation initiation complex eIF4F (Alain et al. 2012; Truitt et al. 2015). This results in the translational repression of mRNAs that are more sensitive to eIF4F levels (Suppl. Fig. 3, Model 1). This makes sense in light of the fact that 4E-BP and eIF4G compete for the same region of eIF4E. However, this sequestration model is paradoxical as it suggests that the weakest eIF4E-binding mRNAs should be the most strongly affected by 4E-BP activity. Yet it is known that translation component-encoding transcripts like ribosomal protein mRNAs are efficiently translated, suggesting that the targets of 4E-BP are normally efficient binders of the canonical translational initiation complex.

This problem has been somewhat ameliorated by the recent identification of LARP1 as a second translational effector, which has been shown to bind directly to the 5’TOP and/or PRTE sequence elements as well as to the 5’Cap (Hong et al. 2017; Lahr et al. 2017). One simple model that follows is that specificity for the 5’TOP and/or PRTE elements comes predominantly from deposphorylated LARP1 and that the binding of the 4E-BP-4E complex to the same mRNA elements and regions is a consequence of this specificity. Otherwise put, the binding of 4E-BP to these elements is in equilibrium with LARP1 binding, which then endows 4E-BP with the same in vivo specificity as LARP1; binding of these proteins to other transcripts and regions is less efficient relative to their association with eIF4F (Suppl. Fig. 3, Model 2).

However, RNA recognition as well as translational inhibition of growth-relevant transcripts is likely to also be a property of 4E-BP, which like LARP1is phosphorylated and regulated by mTOR. In this context, the much stronger repression of Thor targets with dPRTE than targets with dMotif-1 (Fig. 3) is consistent with a functional collaboration of LARP1 and Thor to repress the expression of dPRTE-containing transcripts. We note that the impact of 4E-BP on translation is likely to be even more complicated as it depends on eIF4E expression level relative to that of 4E-BP as well as the status of its multiple phosphorylation sites. Moreover, there are three orthologues of mammalian 4E-BP (1/2/3) with tissue-specific expression.

However, even this model is likely to be oversimplified: LARP1 and the 4E-BP have distinct mRNA targets except for PRET mRNAs (Hong et al. 2017). Moreover, Thor-TRIBE targets containing the dPRTE motif and those with the CGGTCACACT motif (dMotif-1) rarely overlap and therefore probably reflect different groups of regulated targets. This suggests that Thor collaborates with different RBPs that impact and may even determine mRNA recognition. It is therefore possible that THOR has no RNA recognition activity, which appears to conflict with the positive CLIP result (Fig. 4). However, CLIP may be able to reveal intrinsically low affinity RNA-protein interactions. In the case of 4E-BP, tethering to higher affinity RBPs via protein-protein interactions may provide a sufficiently high local concentration near RNA to enable a positive CLIP result. A similar logic may apply to many of the large number of RNA binding proteins revealed by in vivo cross-linking procedures, i.e., perhaps only a minority are RBPs that can bind RNA on their own with high affinity.

It may be relevant in this context that a comparable role of *Drosophila* LARP1 to that of mammalian LARP1 in growth control has not been established. Indeed, there is considerable divergence between the fly and the mammalian protein (data not shown), and *Drosophila* LARP1 is known to have other functions (Zhang et al. 2019). These considerations indicate that even recognition of the dPRTE motif by *Drosophila* LARP1 is currently uncertain.

Although 4E-BP-mediated editing is enriched in the 5’UTR, editing clearly occurs elsewhere within the transcript, in fly cells as well as in mammalian cells. This could reflect 4E-BP overexpression, but it is likely that the ADARcd can also “reach” and edit a susceptible sequence element a considerable distance away from the binding site of its protein complex (McMahon et al. 2016; Xu et al. 2018).

In summary, TRIBE works well for eIF4E-BP, in mammalian cells as well as in the fly system. Recent results from elsewhere on different RNA-binding proteins show that TRIBE works well, in plant (Peter Brodersen, personal communication) as well as in mammalian systems (Biswas et al., 2019; Herzog et al., 2019). Moreover, our current results suggest that TRIBE demonstrates more target transcript specificity than CLIP (data not shown). Although work in this paper does not constitute a proper head-to-head comparison of TRIBE and CLIP, this suggestion is similar to conclusions from other TRIBE vs CLIP comparisons in mammalian cells (Biswas et al., 2019; Biswas personal communication) and may reflect the longer dwell time and therefore tighter binding required for editing than for cross-linking (Kuttan and Bass 2012). At a minimum, all of this progress indicates that TRIBE will add to the arsenal of techniques available to identify and study the mRNA targets of diverse RBPs in many different cells and systems.

## Acknowledgements

We thank Michael Marr, Amy Lee and Katharine C Abruzzi for crucial comments on the manuscript as well as current Rosbash laboratory members for comments and discussion. We also thank Nahum Sonenberg for the pcDNA3-3HA-4E-BP1 plasmid. The work was supported by the Howard Hughes Medical Institute and by a National Institutes of Health grant R01AG052465.

---

**Supplementary Fig. 1.**
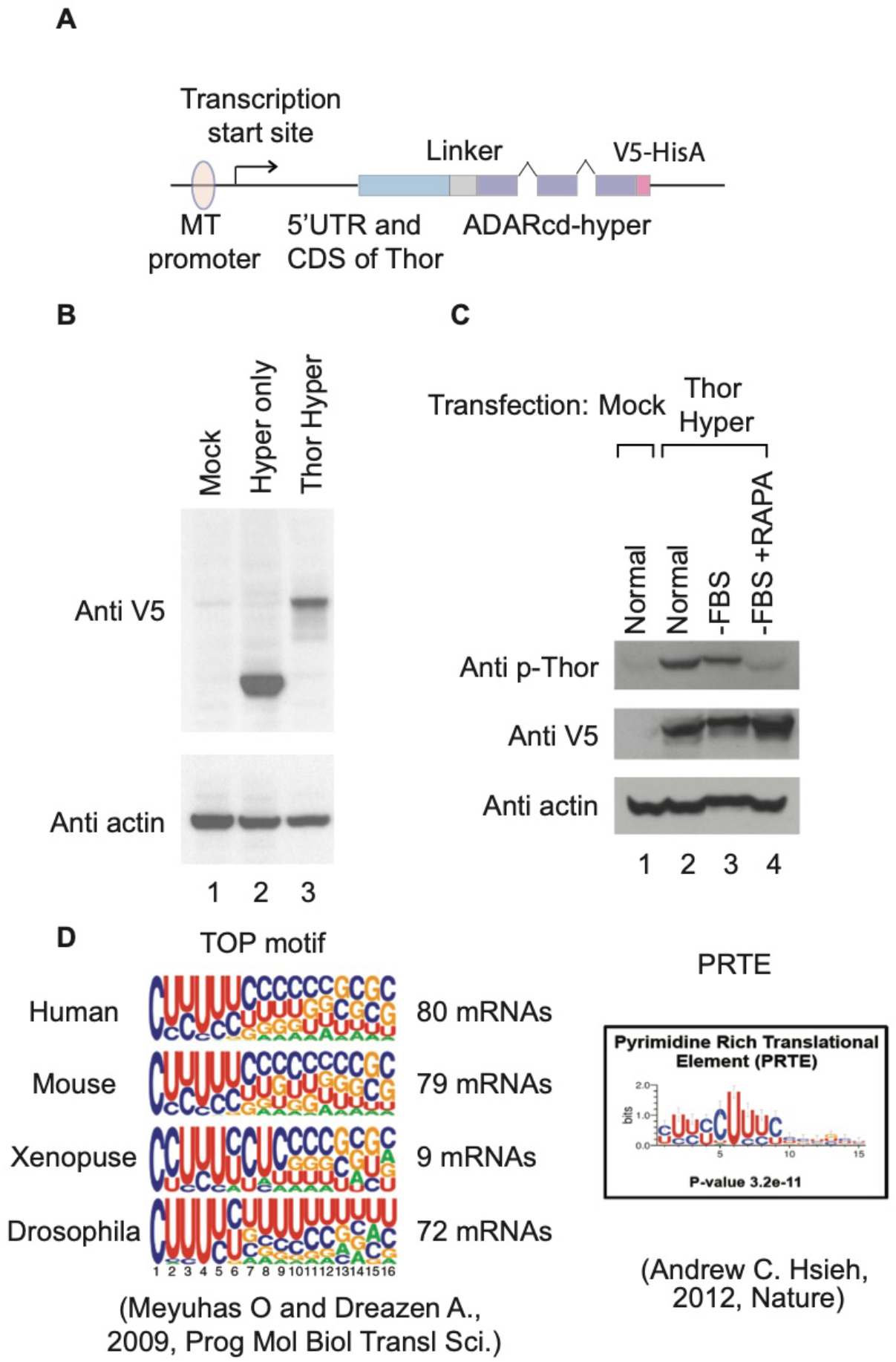
**(A)** Schematic presentation of Thor hyperTRIBE DNA construct. **(B)** The proteins of Hyper (ADARcd-E488Q-V5) and Thor Hyper (Thor-ADARcd-E488Q-V5) are expressed in S2 cells after transfection and induction by copper sulfate. The western blot was carried out using antibody against V5 tag and actin. **(C)** S2 cells were transfect with mock plasmid or the plasmid encoding Thor-ADAR-E488Q-V5 (Thor-Hyper). Cells expressing Thor-ADAR-E488Q-V5 were incubated in media with FBS (Normal), without FBS (−FBS), or with rapamycin in addition to FBS depletion (−FBS +RAPA). Western blot was performed using antibody against phospho-4E-BP1 Thr37/46 (p-Thor), V5 tag, and actin. **(D)** Left: The consensus sequences of 5’ terminal oligopyrimidine tract (5’ TOP) are shown. The TOP motif is conserved in Drosophila rp (ribosomal protein) mRNAs. TOP motif resides from position +1 of the 5’UTR. Right: The consensus sequences of Pyrimidine Rich Translational Element (PRTE) are shown.

**Supplementary Fig. 2.**
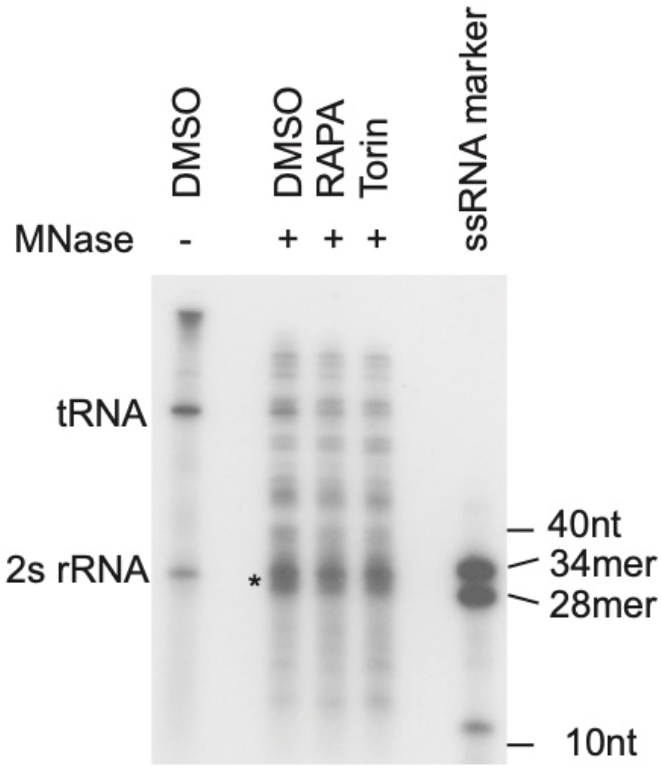
Ribosome protected fragments (RPFs) were enriched between 28-34nt (marked with an asterisk). S2 cells were treated with 100 nM of rapamycin (RAPA), Torin, or DMSO. Cell lysate was treated with MNase and passed through Sephacryl S-400 columns to purify monosomes. RNA was purified from the flowthrough of the columns. 5’-end of RNA was labelled by γ-[32P] ATP, resolved on a 15% TBE-ureagel and visualized.

**Supplementary Fig. 3.**
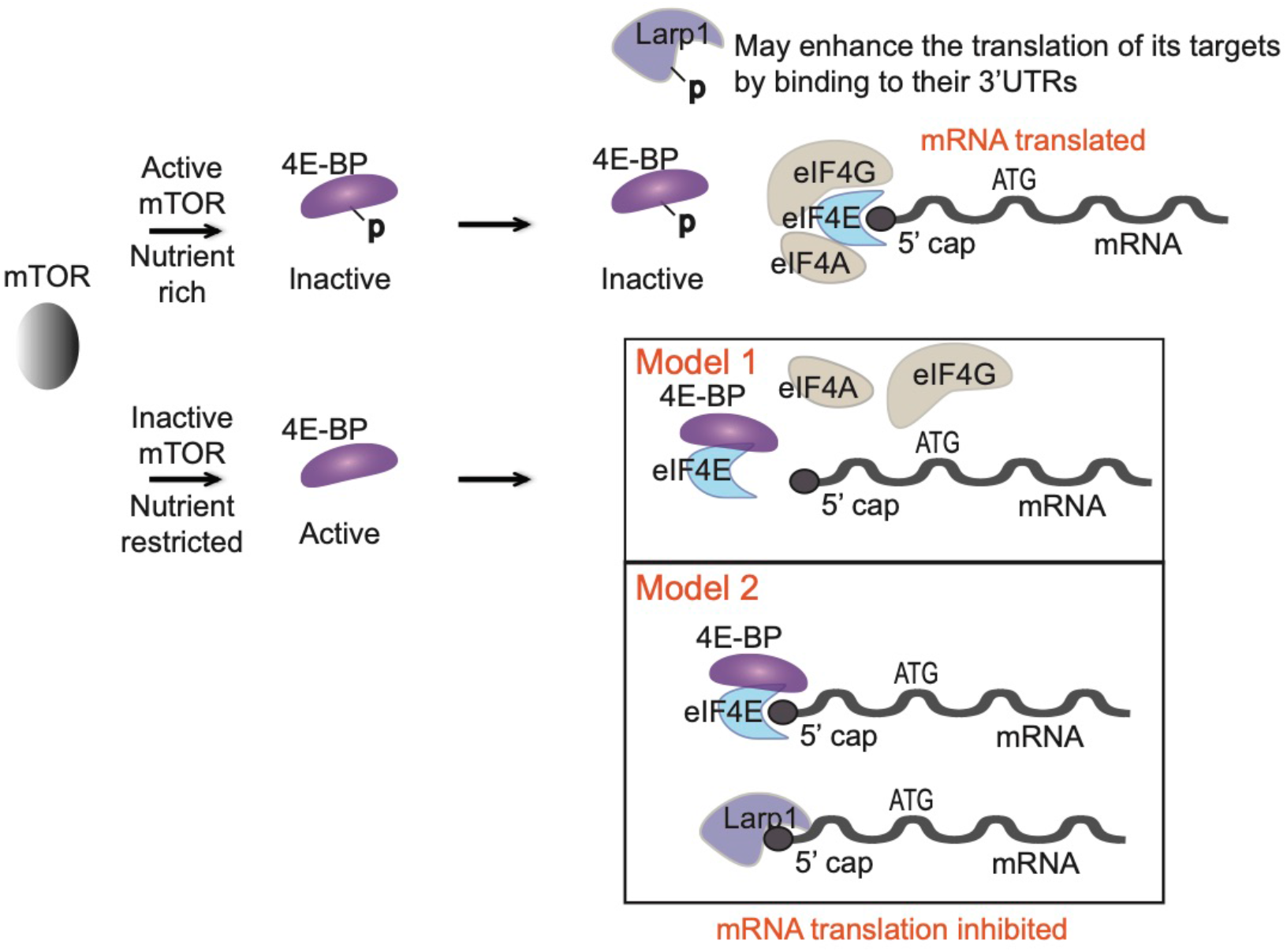
The possible acting mechanism of 4E-BP-mediated translational repression. Under nutrient rich conditions, the mTOR phosphorylates 4E-BP, which prevents it from binding to elF4E. Under nutrient restricted conditions, inactivation of mTOR results in dephosphorylation and 4E-BP activation. Model 1: elF4G and hypophosphorylated 4E-BP share the same binding site on elF4E, active 4E-BP blocks the elF4G binding site on elF4E and inhibits elF4F formation and translation. Model 2: Hypophosphorylated 4E-BP binds to elF4E and subsequently 4E-BP-elF4E complex is associated with their target mRNAs. Both hypophosphorylated Larp1 and 4E-BP target PRTE mRNAs to repress their translation.

## EXPERIMENTAL PROCEDURES

### TRIBE in S2 cells

The region including 5’UTR and CDS of Thor was cloned into pMT-Linker-ADARcd-E488Q-V5 plasmid (100aa flexible linker) to make pMT-Thor-Linker-hyperTRIBE construct using Gibson Assembly (NEB) (Xu et al. 2018). 400 ng of pMT hyperTRIBE plasmid and 400 ng of pAct5.1-eGFP were co-transfected into *Drosophila* S2 cells using Cellfectin II from Thermo Fisher Scientific. 24 hours after transfection, TRIBE protein expression was induced with copper sulfate. Another 24 hours later, cells were treated with 50 nM of rapamycin (Fisher Scientific) plus serum depletion, 50 nM of rapamycin, 50 nM of torin (Fisher Scientific) or vehicle DMSO for 5 hr. More than ten thousand of GFP-positive cells were sorted and collected with a BD FACSAria II machine. Total RNA was extracted from the sorted cells with TRIzol LS reagent. Expression of proteins was assayed by western blot using antibodies against V5 tag (Abcam, ab9116), phospho-4E-BP1 (Thr37/46) (Cell Signaling Tech, # 2855), β-Actin (Santa Cruz Biotechnology, sc-47778).

Standard Illumina TruSeq RNA Library Kit was used to construct an RNA-seq library for TRIBE experiments. FACS-sorted cells were subjected to RNA-seq library protocol as previously described (McMahon et al. 2016). All libraries were sequenced by Illumina NextSeq 500 sequencing system using NextSeq High Output Kit v2 (single-end, 75 cycles). Each sample was covered by ∼10 million raw reads. The TRIBE analyses were followed the process in the paper (Rahman et al. 2018). Briefly, mRNA sequencing reads were mapped to dm3 using Tophat2 (−m 1 -p 5 -g 2 -I 50000 --microexon-search --no-coverage-search) allowing 1 mismatch. PCR duplicates were removed for editing analysis. Only events with minimum 20 reads and 10% editing were considered to be an editing event, gDNA coverage and uniformity of nucleotide identity were also required to avoid inclusion of single nucleotide polymorphisms.

### TRIBE in PC3 cells

The pCMV-hADAR2cd-E488Q plasmid was previously created by making a point mutation on the corresponding position of hADAR2 catalytic domain (Herzog et al., 2019). The region of hADARcd-E488Q was cloned into pcDNA3-3HA-h4E-BP1 (a generous gift from Nahum Sonenberg) by Gibson Assembly to create pcDNA3-3HA-h4E-BP1-hADARcd-E488Q vector. Correct insertion was verified by sanger sequencing.

PC3 cells were obtained from the ATCC and maintained in F-12K Medium (ATCC) with 10% of FBS as suggested by ATCC. pAAV-CMV-eGFP3 was co-transfected into PC3 cells with TRIBE construct using Lipofectamine™ 2000 (Thermo Fisher Scientific). One day or two days later, PC3 cells were treated with 200 nM of INK-128 (MedChemExpress), 2.5 μM PP242 (Selleckchem) or DMSO for 5 hr. GFP-positive cell sorting, RNA-seq library generation, and high-throughput sequencing were carried out as those in S2 cells. The TRIBE analyses were followed the processes in the paper (Rahman et al. 2018). Briefly, mRNA sequencing reads were mapped to hg38 genome using Star3 (--runThreadN 8 --outFilterMismatchNoverLmax 0.07 --outFilterMatchNmin 16 --outFilterMultimapNmax 1). PCR duplicates were removed for editing analysis. The gDNA sequencing data of PC3 were obtained from published data (Seim et al. 2017) to avoid inclusion of single nucleotide polymorphisms. Only events with minimum 20 reads and 10% editing were considered to be an editing events.

### Ribosome profiling in S2 cells

Lysate preparation: The preparation of lysate and ribosome-protected fragment (RPF) for ribosome profiling were generally followed the protocol described in the paper (Dunn et al. 2013). Eighteen million S2 cells were plated to 10 cm-dishes in 12 ml of cell culture media (HyClone SFX-Insect cell culture media with 10% of heat-inactivated FBS [Thermo Fisher Scientific]) one day before experiment. Cells were treated with 100 nM of Rapamycin, Torin1 or vehicle DMSO for 2 hours (6 dishes of cells for each drug). After treating cells with 20 μg/ml of emetine (Sigma-Aldrich, St Louis, Missouri) for 2 min, cells were washed twice with ice-cold PBS (+20 μg/mL emetine) and collected 4–6 cell volumes (200 μl/10 cm dish) of cold polysome lysis buffer (50 mM of pH 7.5 Tris buffer, 150 mM of NaCl, 5 mM of MgCl_2_, 0.5% of Triton x-100, 1 mM of DTT, 20 U/ml of SUPERase• In RNase Inhibitor [Ambion], 20 μg/ml of emetine). Cells were then homogenized on ice in a pre-chilled dounce homogenizer (7 times with A pestle and 7 times with B pestle). The lysate was clarified by spinning 10 min at 20,000 g at 4°C. The supernatant was separated to 3 sets for generating RNA (input) and RPF sequencing libraries. Then 750 μl of Trizol LS (Thermo Fisher Scientific) was added to 250 μl of lysate and total RNA (Input RNA) purification was followed the manufacturer’s instructions. Other sets were flash-frozen in liquid nitrogen, and stored at −80°C for RPF preparation. When ready to proceed, thaw frozen samples on ice in the cold room.

RPF preparation: For each sample, lysate was diluted 2:1 in digestion mixture (50 mM Tris pH 7.5, 5 mM MgCl_2_, 0.5% Triton x-100, 1 mM DTT, 20 U/ml SUPERase• In, 20 μg/ml emetine, 15 mM CaCl_2_, and 150 U/dish micrococcal nuclease [Roche Applied Science]). Samples were digested for 40 min at 25°C in a Thermomixer (Eppendorf, Hamburg, Germany). Digestions were stopped by adding EGTA to a final concentration of 6.25 mM and placing the reactions on ice. For purification of monosomes, Sephacryl S-400 columns (GE Healthcare) were used. Columns were vortexed to mix resin well (avoid bubbles) and the storage buffer on resin was removed by gravity flow. Columns were equilibrated by repeating five times the step of loading 600 μl of polysome buffer and removing it by gravity flow. The columns were spun down at 600G for 4 min. 250 μl of the sample was loaded to the column, spun down at 600G for 2 min to collect the solution flown through into a new tube. The RNA was then purified from the solution using Trizol LS.

rRNA depletion: 10 μg of input RNAs and 2.5 μg of RPF RNAs were treated with Ribo-Zero reaction (MRZH11124C, Epicentre) kit by following the manufacturer’s instructions. For rRNA depletion from RPF RNAs, 50°C incubation what is the step just before collecting rRNA-depleted RNAs in the manufacturer’s instructions was omitted. The collected 100 μl of supernatant from Ribo-Zero reaction is precipitated by adding 500 μl of 100% ethanol, 10 μl of 3M NaOAc, 1 μl of glycoBlue (Invitrogen). The RNA was resuspended in 10 μl of nuclease-free water.

Sequencing library preparation: Input RNA was fragmented by partial alkaline hydrolysis. 10 μl of input RNA was mixed with 10 μl of 2× fragmentation buffer (2 mM EDTA, 12 mM Na_2_CO_3_, 88 mM NaHCO_3_, pH ~9.3) and incubated at 95°C for 18~20 min. Then, input RNA and RPF were resolved on a 15% TBE-urea gel (Invitrogen). A gel slab corresponding to 34–55 nt for input and 28-34 nt for RPF was excised from the gel, eluted, and precipitated. RNA was removed 3’phosphoryl groups using T4 Polynucleotide Kinase (New England Biolabs) in the buffer (70 mM Tris-HCl, 10 mM MgCl_2_, 0.8 mM DTT, PH 6.5) at 37°C for 20 min. After phenol extraction, library preparation was followed the manufacturer’s instructions of TruSeq small RNA kit (Illumina) with modifications. 3’ adaptor (TruSeq small RNA kit, RA3, preadenylated) was ligated to the RNA using T4 RNA ligase 2 truncated (New England Biolabs) by incubating at 25°C for 6 hr, 22°C for 6 hr. 5’end of RNA was phosphorylated and labelled by γ-[32P] ATP using T4 Polynucleotide Kinase (New England Biolabs) at 37°C for 45 min. 3’adaptor-ligated RNA was resolved on a 15% TBE-urea gel (Invitrogen) and the RNA in the size of 54–75 nt for input and 48-54 nt for RPF was purified. 5’adaptor (TruSeq small RNA kit, RA5, a degenerate bar code, four random nucleotides, was added at 3’end of RA5) was ligated to RNA using T4 RNA ligase (New England Biolabs) at 25°C for 6 hr and 22°C for 6 hr. Reverse transcription was carried out using RT primer (TruSeq small RNA kit, RTP) and superscript III RT enzyme (Invitrogen). Library was amplified by 10–14 cycles of PCR using indexing primers (TruSeq small RNA kit) and phusion polymerase (Thermo Fisher Scientific). Amplification product was size-selected on 6% TBE gels (Invitrogen). Samples were then quantitated using the Bioanalyzer High Sensitivity DNA assay (Agilent Technologies), and single-end sequenced in Illumina NextSeq 500 sequencing system using NextSeq High Output Kit v2 (75 cycles).

### Sequence processing and alignment of ribosome profiling

The alignment of sequencing reads was followed the step 90-96 in the paper (Moore et al. 2014). Briefly, the law reads were filtered and pre-processed according to quality scores (fastq_filter), collapsed the exact sequence duplicates (fasta2collapse), trimmed 3’adaptor sequence (fastx_clipper), remove degenerate bar-code sequences (stripBarcode). The reads were aligned to the reference genome dm3 (novoalign -t 85 -l 23 -s 1 -r None) at least 23 high-quality matches and with alignment cost score ‘−t 85’. Then, collapsed the potential PCR duplicates by coordinates and identified unique reads.

mRNA abundance (input RNA) and ribosome density (RPF) for each genomic feature were measured in fragments per kilobase of feature length per million reads aligning to genomes (FPKM). mRNA abundance reflects the total number of RNA fragments aligning to all countable exonic positions for a given gene. Ribosome density means RPF fragments aligning to all countable positions of a coding region (CDS) for a given gene. We calculate translation efficiency (TE) as the ratio of footprint (RPF) FPKM in the CDS to the RNA fragment (Input RNA) FPKM across the exons. FPKM more than 2 was used for further analysis.

### Motif analysis and gene ontology

We performed motif analysis using MEME version 4.11.2 (Bailey et al. 2009) (parameters: -minw 6 -maxw 10 -maxsize 10000000 -dna -nmotifs 5 -maxsites 200). Gene ontology analyses were performed using PANTHER (Mi et al. 2019). TRIBE target genes were analyzed against a background of all genes expressed in the S2 cells or PC3 cells (>2 fpkm). Gene expression levels were quantified using Cufflinks2.

