## Supplemental Tables for "TRIBE editing reveals specific mRNA targets of eIF4E-BP in *Drosophila* and in mammals": Suppl table 3-Thor TRIBE targets reduced in RP.docx

One hundred forty-four genes are detected as thor targets in thor Hyper-TRIBE and also the translational efficiency of them are decreased after treatment of mTOR inhibitor.

(Thor Hyper-TRIBE: Rapa_common + Torin_common

Ribosome profiling: TE<0.8, p<0.1)

**Ribosome proteins (39)：**

RPL5, RPL7, RPL7-LIKE, RPL9, RPL10, RPL10AB, RPL15, RPL18A, RPL23A, RPL24-LIKE, RPL26, RPL27, RPL28, RPL29, RPL31, RPL32, RPL35, RPL36A, RPL37A, RPL38, RPL41, RPLP0, RPLP1, RPLP2, RPS11, RPS13, RPS14B, RPS15AB, RPS16, RPS18, RPS26, RPS27, RPS27A, RPS28B, RPS29, RPS3A, RPS6, RPS7, mRPS9

**Translational initiation factors (11)**

**eIF3b (eIF3-S9), eIF3d1 (eIF-3p66), eIF3e (Int6), eIF3g1 (CG8636), eIF3h, eIF3i (Trip1), eIF3j, eIF3k (CG10306), eIF3l (CG5642), eIF3m (Tango7)**

eIF-2alpha

**Other translation related proteins (7)**

EEF1DELTA, EF1ALPHA48D, EF1GAMMA, ERF1, EIF-5A, AATS-TYR, NOP5(rRNA modification)

**Other proteins (87)**

ZFRP8

TCTP

SRP54K

SRP19

SLMO

SHRB

SESB

SAM-S

RPII33

RPB10

RPA-70

RM62

RHEB

REG

RCD5

RACK1

RAB1

PROSALPHA7

PROSALPHA5

POLO

PIX

PCMT

PCH2

NLP

NAP1

Mpcp2

MGR

MDH1

LSM7

LOST

L(3)01239

KARYBETA3

JHI-26

IPK2

HSC70-5

GAPDH1

FDH

ENDOGI

EFF

EDL

DROJ2

COVB

COOP

CG9836

CG9350

CG9330

CG8386

CG8353

CG8258

CG7789

CG7265

CG6686

CG6523

CG6388

CG6353

CG6195

CG5854

CG5355

CG44014

CG34179

CG34164

CG32267

CG31548

CG31436

CG30273

CG18343

CG16985

CG16817

CG15735

CG1550

CG15083

CG13630

CG13319

CG1236

CG12173

CG11752

CG11454

CG10565

CCHL

BEAF-32

BALL

ATPSYN-B

ARPC2

ARF79F

AOS1

ANT2

ADK2
